# Resilience to multiple stressors in an aquatic plant and its microbiome

**DOI:** 10.1101/726653

**Authors:** Anna M. O’Brien, Zhu Hao Yu, Dian-ya Luo, Jason Laurich, Elodie Passeport, Megan Frederickson

## Abstract

**Premise:** Environments are changing rapidly, and outcomes of species interactions, especially mutualisms, are notoriously dependent on the environment. A growing number of studies have investigated responses of mutualisms to anthropogenic changes, yet most studies have focused on nutrient pollution or climate change, and tested single stressors. Relatively little is known about impacts of simultaneous chemical contaminants, which may differ fundamentally from nutrient or climate stressors, and are especially widespread in aquatic habitats.

**Methods:** We investigated the impacts of two common contaminants on interactions between the common duckweed *Lemna minor* and its microbiome. Sodium chloride (salt) and benzotriazole (a corrosion inhibitor) negatively affect aquatic organisms individually, yet commonly co-occur in runoff to duckweed-inhabited sites. We tested three *L. minor* genotypes with and without the culturable portion of their microbiome across field realistic gradients of salt (3 levels) and benzotriazole (4 levels) in a fully factorial experiment (72 treatments), and measured plant and microbial growth.

**Key Results:** We found that stressors had conditional effects. Salt decreased both plant and microbial growth, but decreased plant survival more as benzotriazole concentrations increased. In contrast, benzotriazole did not affect microbial abundance, and benefited plants when salt and microbes were absent, perhaps due to the biotransformation we observed without salt. Microbes did not ameliorate duckweed stressors, as microbial inoculation increased plant growth, but not at high salt concentrations.

**Conclusions:** Our results suggest that multistressor effects matter when predicting responses of mutualisms to global change, but that mutualisms may not buffer organisms from stressors.

## INTRODUCTION

Human activities are exposing many organisms to new or intensifying stressors, with effects often cascading to other community members via species interactions. Mutualisms often link the fates of interacting species, and most organisms engage in at least one mutualism with another species, be it for nutrition, protection, or dispersal. There is concern that mutualists may be particularly vulnerable to environmental change because mutualism outcomes are often context-dependent. In other words, the magnitude of benefits exchanged between mutualistic partners typically depends on the external abiotic or biotic environment (Chamberlain et al., 2014), and environments are changing at a historically unprecedented pace. As a result, many mutualisms are now taking place in novel environments where partners face multiple stressors (Kiers et al., 2015). A key question is therefore: how do anthropogenic stressors, singly or in combination, affect mutualism outcomes? And in the same vein, does mutualism mitigate or exacerbate global change (Frederickson, 2017)?

There is also currently substantial interest in plant-associated microbiomes and how they may facilitate acclimation or adaptation to plant stressors. The term ‘microbiome’ refers either to the community of microbes living together in an environment, such as on or in a host, or their collective genomes. Many plant-associated microbes are beneficial, or at least not pathogenic, and there is growing appreciation for their effects on plant physiology, ecology, and even evolution (Lau and Lennon, 2012; Panke-Buisse et al., 2015). Much of what we already know about context-dependent outcomes in mutualisms comes from studies of plant rhizosphere microbes, especially legume-rhizobium and plant-mycorrhizae interactions in terrestrial ecosystems (Heath and Tiffin, 2007; Hoeksema et al., 2010). A number of recent studies have evaluated the effects of stressors such as nutrient pollution or climate change on these interactions (Shantz et al., 2016). For example, Weese et al. (2015) found that 22 years of nitrogen deposition caused the evolution of less beneficial rhizobia, suggesting that anthropogenic nitrogen inputs to ecosystems may weaken legume-rhizobium mutualisms. More generally, a meta-analysis by Shantz et al. (2016), which included both plant-microbe and other mutualisms between heterotrophs and autotrophs (e.g., corals), found that nutrient loading decoupled the performance of mutualistic partners, with autotrophs benefiting at the expense of heterotrophs.

Most real ecological communities face multiple threats, as climate, nutrients, and contaminants all change in tandem, and feedback loops connect abiotic variables to shifts in community structure. Even amidst widespread recognition of this complexity, research still focuses on how single stressors affect plant-microbe mutualisms, because studying multiple stressors remains challenging. A key hurdle when assessing the potentially interactive effects of multiple stressors on the outcome of any species interaction is the highly multifactorial nature of the necessary study design. To overcome this hurdle, we have recently developed a novel experimental system to test the effects of multiple stressors on plant-microbe interactions. We use one of the world’s smallest angiosperms, *Lemna minor* (here, duckweed), and its microbiota to simultaneously test many combinations of environmental factors across host genotypes and microbial communities.

Much like terrestrial plants, *L. minor* associates with diverse microbial communities containing beneficial bacteria such as the nitrogen-fixer *Ensifer adhaerens* and plant growth-promoting *Pseudomonas* spp., among others (Ishizawa et al. 2017 and see Results, below). *Lemna minor* mostly reproduces clonally, producing new plants by budding, although in the wild it also occasionally makes (tiny) flowers and set seeds (Landolt, 1975). Clonal populations of *L. minor* grow very fast, allowing us to measure meaningful changes in population growth rate in as little as 3 days (e.g. Liu et al., 2017). *Lemna minor* occurs at high densities in slow-moving fresh water under a wide range of environmental conditions and in many parts of the world (Landolt, 1975; RDSC, 2016), with some populations regularly exposed to aquatic contaminants. There is interest in using *L. minor* or other duckweeds for bioremediation of contaminated water (Mo et al., 1989; Stout et al., 2010; Stout and Nüsslein, 2010; Sekomo et al., 2012; Gatidou et al., 2017), and microbes may amplify or underlie a number of duckweed bioremediation effects (Toyama et al., 2009; Ogata et al., 2013), so understanding how duckweed-microbe interactions respond to multiple chemical stressors has potential real-world implications. Studies have suggested that we might be able to engineer plant microbiomes for particular applications (Mueller and Sachs, 2015), including water treatment using duckweed-microbe associations (Zhao et al., 2015), but to do so we need to understand interactions among plants, microbes, and contaminants. Here, we examine the main and interactive effects of two commonly co-occurring chemical contaminants in urban freshwater, sodium chloride (NaCl or salt) and benzotriazole (BZT), on the outcome of duckweed-microbe interactions. In many northern cities where duckweed is common, large amounts of salt are applied to roads and other surfaces in winter; the city of Toronto alone applies 130,000 - 150,000 tonnes of road salt per year, not including private and residential applications (Transportation Services Division, 2016). The widespread use of salt for road de-icing significantly increases the salinity of urban lakes and ponds (Novotny et al., 2008), and many North American lakes are now at risk of salinization past the U.S. EPA aquatic life threshold criterion (Dugan et al., 2017).

BZT is a corrosion inhibitor widely used in fluids that come in contact with metals, including deicing fluids and engine coolants as well as household products such as dish soaps (Davis et al., 1977; Cancilla et al., 1997; Pillard et al., 2001; Giger et al., 2006; Vetter and Lorenz, 2013). Like salt, BZT contamination of water bodies peaks in the late winter or early spring (Alvey et al., 2016; Parajulee et al., 2017), possibly due to increases in BZT use in antifreezes and de-icers for cars, airplanes, or other heavy equipment (Kiss and Fries, 2012). Existing evidence suggests that BZT harms aquatic plants and animals, including duckweed (Pillard et al., 2001; Seeland et al., 2012; Gatidou et al., 2017), and that it may be a human carcinogen (Richardson and Ternes, 2018). BZT resists degradation in the aquatic environment (Weiss et al., 2006), but green infrastructure methods may remove substantial amounts via biodegradation or sorption (e.g. Matamoros et al., 2010; Rhodes-Dicker and Passeport, 2019), for which duckweed shows initial promise (Gatidou et al., 2017).

In a full factorial experiment, we tested the main and interactive effects of three NaCl concentrations, four BZT concentrations, and the presence or absence of microbes on the growth of three clonal genotypes of the duckweed *L. minor*. Specifically, we asked whether NaCl or BZT reduces plant or microbial performance, and at what concentrations. We also investigated whether NaCl and BZT have non-additive effects on plants and microbes, and whether microbes ameliorate or exacerbate the effects of these potential stressors on plants. Finally, we investigated whether plants, microbes, or both break down BZT under our experimental conditions, raising the possibility that they could potentially be used for bioremediation of this emerging contaminant.

## METHODS

### Biological materials

We collected many *Lemna minor* plants from each of three water bodies in the Greater Toronto Area during summer 2017, and separately cultured the plants and their associated microbes (BH = Bruce Handscombe Memorial Park, KSR = Koffler Science Reserve, MT = Moccasin Trail Park; exact site locations in Table S1). We cultured plants and microbes from freshly collected live tissue from each site, but we also froze some *L. minor* plants from each site for molecular work (see below). *Lemna minor* is a small aquatic angiosperm that can reach high densities in ponds and slow-moving streams. It has a nearly cosmopolitan distribution (RDSC, 2016), and is widespread in the Greater Toronto Area, including in stormwater ponds and other water bodies that are commonly exposed to runoff. Duckweeds have microbial communities living on or in their roots and fronds, and our previous work has found that experimental microbial inoculation of sterilized *L. minor* increases plant growth (O’Brien et al., 2018).

We cultured isogenic *L. minor* lines by transferring one field-collected live frond to growth media (recipe in Krazčič et al. 1995). In approximately three months, cultures grew from this single frond to high densities (~500 plants per 500 mL jar) of isogenic individuals that we used in experiments. We grew and maintained these lines in 500 mL vented glass jars in a growth chamber with a cycle of 16 hours at 23°C and 150 *μ*mol/m^2^ and 8 hours of 18°C in the dark. *Lemna minor* can flower and set seed in the field, but genetic evidence suggests it does so only very rarely (Ho, 2017). We never observed flowers in our cultures, suggesting very little or no recombination occurred under our growth conditions in the lab. Furthermore, *L. minor* has very little segregating diversity within ponds to begin with (Ho, 2017), and as such, even with recombination, the difference between our isoparental lines and a definitively isogenic line should be negligible.

To extract microbes from field-collected *L. minor*, we pulverized freshly collected tissue of one or two fronds from each site. We then swabbed pulverized tissue onto yeast mannitol media (YMA) agar plates, cultured them at 29°C for 5 days, and stored them at 4°C until use in experiments. The cultured microbes thus comprise the subset of all microbes associated with *L. minor* that can grow on yeast mannitol media.

### Sequencing of microbial communities

We used 16S rDNA amplicon sequencing to characterize the bacterial communities both in our field samples of *L. minor* and in the microbial cultures we used in our experiments. We extracted DNA directly from frozen field-collected *L. minor* tissue using DNeasy Powersoil (Qiagen) kits. We extracted DNA from our yeast mannitol media cultures using GenElute Bacterial Genomic DNA (Millipore-Sigma). We sent 10 ng of DNA of each field and culture sample to Genome Québec (McGill University) for PCR amplification of 16s rDNA, normalization, and barcoded, paired-end, 250 base pair format sequencing on an Illumina MiSeq System. We sequenced the V3-V4 variable region with primers 341f - 805r, as this region has good recovery and little bias for plant microbiomes (Thijs et al., 2017). We received demultiplexed reads from the facility (specifically 62,355, 119,337, and 107,759 reads for BH, KSR, and MT field samples, and 77,294, 103,451, and 110,183 reads for BH, KSR, and MT cultures, respectively) totalling 290,189,500 base pairs. We processed reads with QIIME2 (Bolyen et al., 2018), using deblur to correct errors and to assign amplicon sequence variants (ASVs), the naive Bayesian classify-sklean pre-trained on the Greengenes database (McDonald et al., 2012) to assign taxonomy to ASVs with a confidence of 70% or higher, and filter-taxa and filter-table to remove chloroplast reads. We summarized data in R and used package ade4 for principal components analysis (Dray and Dufour, 2007; R Core Team, 2014).

### Chemicals

Benzotriazole (BZT, purity = 99%) and hexane (purity ≥ 98.5%) were purchased from Sigma-Aldrich (Oakville, Canada). Ethyl acetate (purity ≥ 99.8%) and methanol (purity ≥ 99.9%) were obtained from EMD Millipore (Etobicoke, Canada), and Fisher Scientific (Whitby, Canada), respectively. Hydrochloric acid (36.5% to 38.0% solution in water) was purchased from Caledon Laboratories (Georgetown, Canada). Milli-Q (18.2 MΩcm) water was supplied by a Millipore ultra-pure water system (Etobicoke, Canada). Sodium chloride (NaCl, purity ≥ 99.5%) was purchased from Sigma-Aldrich (St. Louis, Missouri).

### Experimental design and data collection

We chose three concentrations of NaCl and four concentrations of BZT to add to a base media solution (Krazčič et al., 1995) in a factorial design. The media itself had negligible salt (Krazčič et al., 1995) and BZT concentrations (verified by HPLC/DAD as described below, data not shown). Either no salt or 0.8 or 10 g/L of salt as NaCl was added to the media. The 10 g/L level is near the reported maximum tolerance of *L. minor* to salt (Sree et al., 2015; Liu et al., 2017), while 0.8 g/L is a representative salt concentration for Ontario urban runoff (Rhodes-Dicker and Passeport, 2019). We then crossed these salt treatments with BZT treatments by adding no BZT, 0.1 mg/L, 1 mg/L, or 10 mg/L BZT. Concentrations near 0.1 mg/L are commonly observed for BZT, while 10 mg/L BZT is a high concentration that is very rarely observed in aquatic environments (Loos et al., 2010; Parajulee et al., 2017). We prepared the various experimental solutions by autoclaving salt and media solutions, then mixing with sterile (verified by swabbing on YMA agar and incubating at 30°C) BZT solutions.

We crossed the 12 combinations of salt and BZT concentrations with two microbial treatments: a treatment in which no microbes were added and a treatment in which we inoculated plants with microbes cultured from the same field site. In all treatments, plants were first sterilized in 0.5% bleach for 30 seconds to remove microbes. We prepared microbial inocula from each site by swabbing across the stored agar plate, adding to liquid YMA, culturing in a shaker for 3 days at 30 °C and 200 rpm, and diluting to control cell density across inocula. We diluted by measuring optical density, converting to cell counts by applying a single calibration curve across inocula (correlated to colony forming units, O’Brien et al., 2018), and standardizing to approximately 3000 cells per *μ*L. The exact relationship of optical density to cell count depends on both cell size and composition (Volkmer and Heinemann, 2011), both of which likely vary across the taxa in our cultures, so cell concentrations may differ slightly across inocula.

Each of the three source populations of duckweed received all 24 salt-BZT-microbe treatments, for a total of 72 treatments. Each treatment was replicated eight times, for a total of 576 experimental units. We grew the experiment in 24-well plates, in a completely randomized design. Each well received 2.5 mL of one of the treatment solutions, one cluster of duckweed fronds selected from cultures without regard to size (generally one mother-daughter frond pair), and 20*μ*L of microbial inoculum or 20*μ*L of sterile microbial medium. We sealed plates with BreatheEasy membranes (Millipore-Sigma) to prevent microbial cross-contamination between wells or from outside the plate while allowing gas exchange. We put the 24 well plates in the growth chamber under the same conditions as duckweed cultures, and let plants and microbes grow for 10 days.

We photographed plates immediately after setup (day 0), and again on days 4, 7, and 10. Using two custom-built camera rigs, photographs were taken under two lighting regimes and camera distances, one from above for days 0 and 10, and one from below for days 4 and 7, when plate seals prevented imaging from above. After standardizing by well diameter, size measurements are comparable across both plates and time points, and all other measures are comparable across plates within time points. We processed images in ImageJ (Schneider et al., 2012) to calculate the pixel area of live duckweed in each well at each time point. If a well had no pixels passing ImageJ thresholds, we regarded the duckweed as dead. Because duckweed primarily reproduces asexually, the pixel area of live fronds is very tightly correlated to the number of reproductive individuals (O’Brien et al., 2018). From images taken on day 10, we also scored two plant traits: greenness and aggregation (the ratio of frond area to frond perimeter, see Figure S1). Greenness is expected to correlate with chlorophyll content (Adamsen et al., 1999; Keenan et al., 2014). The aggregation of duckweed fronds may increase shading in ponds, potentially modulating water temperature, and is therefore a trait of interest. Warm outflow water from stormwater ponds can raise stream temperatures, potentially harming fish (Chu et al., 2005; Herb et al., 2009; Comte et al., 2013).

After photographing plates on day 10, we immediately froze them at −20°C and stored them frozen until we sampled well solutions to measure optical density and BZT concentration (below). We took two 70*μ*L samples from each well, and measured optical density at 600 nm wavelength on a plate reader (BioTek Synergy HT with Gen5 1.10 software). Optical density at 600 nm is proportional to the density of all microbial cells in the sample (O’Brien et al., 2018); i.e., it is a measure of total microbial abundance.

### Evaluation of benzotriazole transformation

We further analyzed a subset of samples to measure the reduction in BZT concentration that occurred by the end of the experiment. Sample plates at −20°C (stored in a freezer for less than 4 months) were thawed. It was not feasible to test all 576 well solutions, so we chose to analyze the BZT concentration at the end of the experiment at the two levels of NaCl with the most different growth outcomes (0 and 10 g/L, see below) and the two most different starting concentrations of BZT that were greater than 0 mg/L (0.1 and 10 mg/L). Thus, for each of the three genotypes, we selected three replicates at random from the eight possible for each combination of microbial treatment (inoculated or not), NaCl treatment (0 or 10 g/L), and BZT treatment (0.1 or 10 mg/L), totalling 72 wells. To compensate for water evaporation during the experiment, each replicate was brought back to the initial volume of 2.5 mL using reverse osmosis water. Three replicate wells within each test condition were then pooled by genotype to give a combined sample, reducing 72 samples to 24. Pooling replicate wells within a treatment was required to generate a large enough sample volume for analysis. Pooled samples with an initial benzotriazole concentration of 10 mg/L were directly injected into a Dionex UltiMate 3000 Series high-performance liquid chromatography system equipped with a diode-array detector (HPLC/DAD, Fisher Scientific, Ottawa, Canada) for concentration quantification. The HPLC/DAD was further equipped with an Accucore C18 column (100 × 2.1 mm × 2.6 *μ*m, Fisher Scientific, Ottawa, Canada).

All samples were first filtered with a 0.22 *μ*m nylon filter (Chromatographic Specialties, Brockville, Canada), then analyzed at 30°C using an isocratic elution of 70:30 v/v Milli-Q water and methanol, at a flow rate of 0.1 mL/min and a detection wavelength of 275 nm. For pooled samples with an initial concentration of 0.1 mg/L, solid phase extraction (SPE) was conducted prior to the HPLC/DAD analysis to increase the detection signal above the HPLC/DAD limit of detection (0.1 mg/L). The samples were first acidified to pH 2 by using hydrochloric acid at 1 M. The SPE cartridges (Oasis HLB, 6 mL, 200 mg, Milford, USA) were conditioned successively with 15 mL of hexane, 10 mL of ethyl acetate, 5 mL of methanol, and 10 mL of acidified Milli-Q water at pH 2. Subsequently, the samples were percolated through the cartridges with a flow of 1 mL/min. Then, the cartridges were washed with 5 mL acidified Milli-Q water at pH 2, dried under vacuum for 20 minutes, and eluted with 5 mL of ethyl acetate. The extracts were then dried under a gentle stream of nitrogen and reconstituted in 1 mL of Milli-Q water. Finally, the reconstituted extracts were analyzed for the concentration of benzotriazole using the HPLC/DAD procedure outlined above.

### Data analysis

We analyzed data in R (R Core Team, 2014). Using the MCMCglmm package (Hadfield, 2010), we fit separate linear models to duckweed survival, pixel area over time (excluding wells in which all duckweed died), as well as duckweed traits, optical density (inoculated wells only), and BZT concentration at the end of the experiment. Models included the fixed main and interactive effects of salt, BZT, and inoculation with microbes, as well as a random intercept for each duckweed genotype. For the model of duckweed pixel area, we also included a random intercept for each individual well to account for repeated measures in time. Effectively, this means that treatments were fit as adjustments to the slope of the relationship between pixel area and time (i.e., days).

Growth in pixel area of surviving plants was normally distributed at 0 and 0.8 g/L NaCl, but growth at 10 g/L NaCl was highly right-skewed (Figure S2). Thus, we separately modeled growth at each treatment level of NaCl with a Gaussian or Poisson distribution, as appropriate. We natural log-transformed optical density (plus our approximate detection limit of 0.01 so that values remained finite) to meet normality assumptions. We tested the effects of BZT and salt on optical density only in the inoculated treatment, but we also verified the effect of inoculation on optical density using the full dataset in a separate linear model with inoculation as the only fixed effect and source population as a random effect.

For each response variable, we fit the full model with all two- and three-way interaction terms first. However, since non-significant interaction terms can hide significant component parts, when highest order terms were not significant, we removed them and fit simpler models, repeating if necessary until all non-significant interaction effects were removed. Finally, we removed all non-significant fixed effects, unless they were components of significant interaction terms, and re-fit. For the model of plant growth, we used deviance information criteria (Spiegelhalter et al., 2002) to choose the best-fitting model between models with interaction terms for days or days squared.

Since we measured final benzotriazole concentrations on a subset of the data (24 pooled samples representing 72 wells, see above), we included only treatment main effects of salt, inoculation, and BZT level, with plant genotype again as a random effect. We also tested if remaining benzotriazole was correlated to duckweed or microbial growth in separate fixed-effect models, and in a model with both duckweed and microbial growth included, again with plant genotype as a random effect. We fit all models across response variables with 100,000 iterations, 2,000 burn in, thinning by 100 for Gaussian and Poisson distributed response variables. We increased to 1,000,000 iterations for binomially distributed responses.

## RESULTS

### Microbial community composition

As expected, the culture samples had fewer ASVs than the field samples, and differed in taxonomic composition. Gammaproteobacteria dominated the cultured microbiota, while field samples of *L. minor* contained both Gammaproteobacteria and many other bacterial taxa, especially Alphaproteobacteria and Betaproteobacteria, that were not well represented in cultures (Figure 1a). We detected 70 ASVs in the BH field sample (14 in BH culture sample), but many more from KSR and MT samples (696 and 802 from field, respectively, and 17 and 38 from cultures). Many ASVs were unique to individual samples (1,209 out of 1,410), especially field-collected samples. Furthermore, some ASVs overlapped between field and culture samples from the same site (1,5, and 13 from BH, KSR, and MT), but most field taxa were not cultured (Figure 1a). BH cultures had the most ASVs that did not overlap with their paired field sample, perhaps because of lower coverage. Differences between field and culture samples could also have resulted from PCR biases in amplification or real within-site variation in the microbiota of plants: we used different plants for microbial culturing and 16S sequencing. Nonetheless, when we consider the genus-level identification of ASVs (though not all taxa were assigned a genus), only three genera were not shared between culture samples and their corresponding field samples, and two of those were present at another field site. Likewise, all cultured bacterial families overlapped with families found at at least one field site (Figure 1b). Finally, the principal components analysis (PCA) of families found in inocula revealed some similarity along PC1 between field and inocula communities from the same location (Figure 1c). We did not conduct PCAs at lower taxonomic levels because matrices become very sparse.

**Figure 1:**
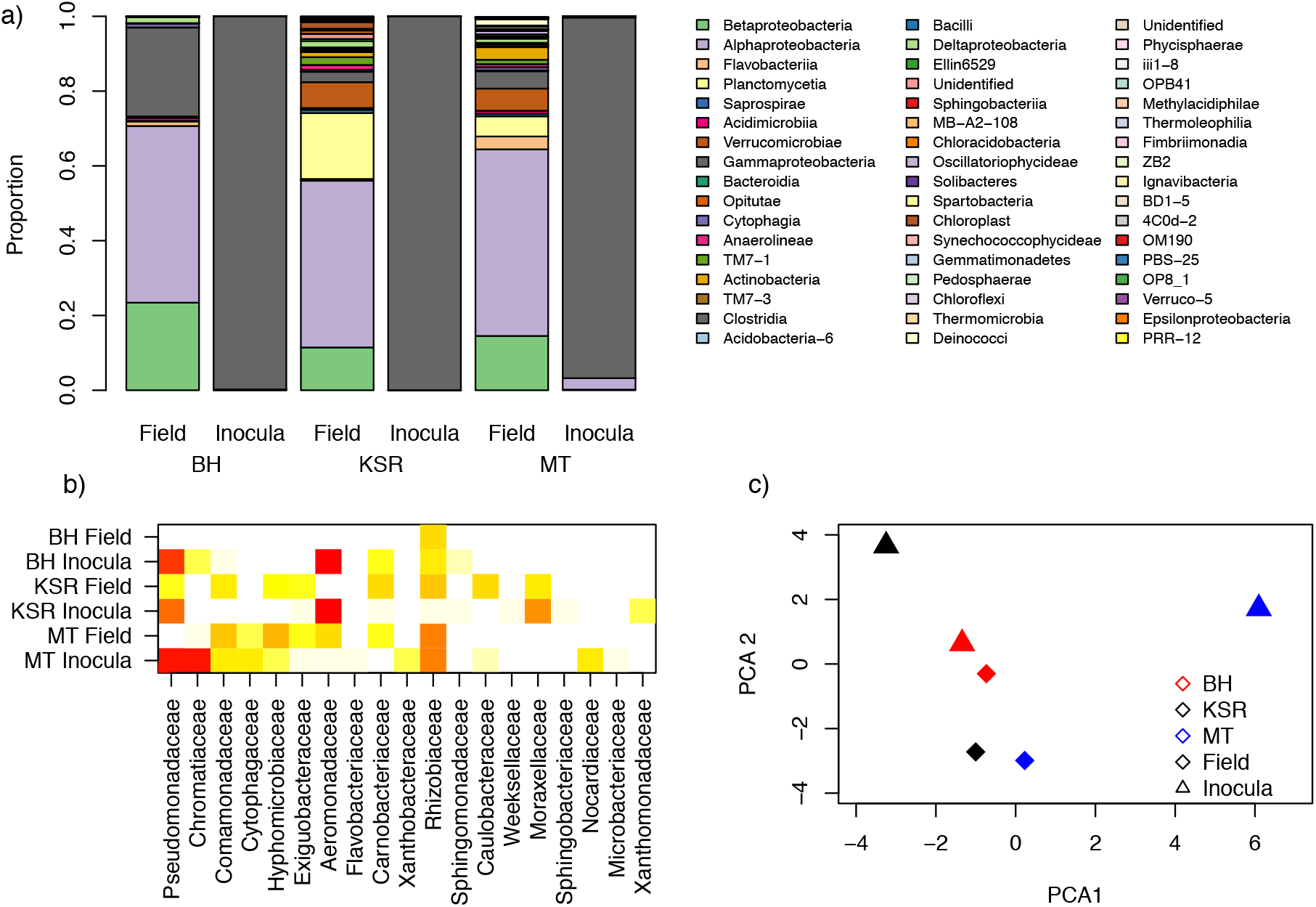
Field and culture-based samples (i.e., inocula) of *L. minor*’s microbiome. a) Bacteria in each sample type at each site (BH = Bruce Handscombe Memorial Park, KSR = Koffler Science Reserve, MT = Moccasin Trail Park) identified to class based on 16s rRNA sequences. Bars show proportions of reads in each class. b) Heatmap of logged proportional abundance of bacterial families in each sample type at each site, subset to families present in cultured samples only. Redder indicates higher relative abundance. c) PCA plot of the data in b), with sites distinguished by color and sample type by shapes.

### Plant fitness through time

The two strongest factors influencing plant growth were NaCl concentration and inoculation treatment. Increasing NaCl concentration generally reduced plant growth and inoculation with microbes generally increased plant growth, however the effect of inoculation treatment depended on NaCl concentration (Table 1). Over the ten days of the experiment, plants grew at 0 and 0.8 g/L NaCl, but high NaCl (10 g/L) concentration resulted in tissue death followed by a modest recovery (Figure S3, Table 1). At high salt, inoculation with microbes slightly reduced growth (Table 1), although this affected mainly growth curves: plant final size was similar between inoculated and uninoculated wells at high salt (Figure 2) and 95% highest posterior density intervals (HPDIs) for predicted mean size overlapped at day 10. In all other treatments, inoculation primarily increased growth (Figure 2, Table 1).

**Table 1:** Model results for plant fitness measures. Regressions for pixel area at 10 g/L NaCl were performed on the log-link scale and parameter estimates are not back-transformed (see Figures 2 and S3 for effect sizes). Significance codes:. pMCMC < 0.1, * pMCMC < 0.05, ** pMCMC < 0.01.

| parameter | Salt concentration (NaCl g/L) |  |  |
| --- | --- | --- | --- |
|  | 0 | 0.8 | 10 |
| Intercept | 1693* | 1701** | 7.33** |
| Days | 28.32 | -24.32 | -1.08** |
| Days<sup>2</sup> | 13.19** | 15.55** | 0.095** |
| Days<sup>2</sup>:BZT | 0.66* | n.s. | n.s. |
| Days<sup>2</sup>:Inoc | 11.62** | 13.68** | -0.0083* |
| Days<sup>2</sup>:BZT:Inoc | -0.79. | n.s. | n.s. |

**Figure 2:**
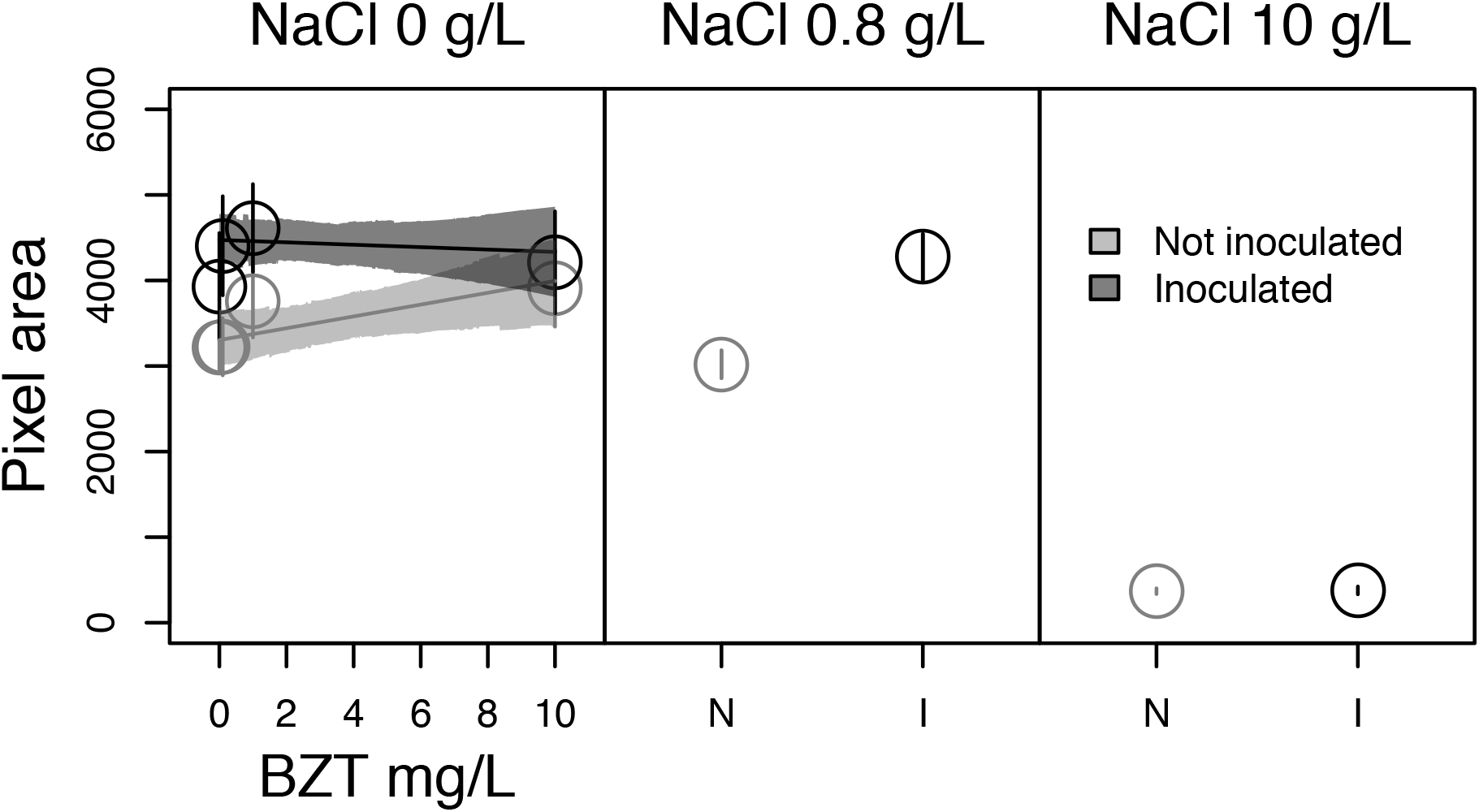
Pixel area of surviving plants at day 10. Each NaCl treatment was analyzed separately. Points are averaged for treatments with significant effects in models, pooling across non-significant treatments (see Table 1). Bars represent one standard error. Model predictions for BZT effects are shown for 0 g/L NaCl data only (predicted mean lines and 95% HPDIs); BZT effects were not significant at higher NaCl concentrations. Microbial inoculation (I on x-axis) is indicated by dark gray circles and intervals; uninoculated wells (N on x-axis) are shown with light gray circles and intervals.

The effect BZT on plant growth depended on NaCl concentration and the presence of microbes. At 0 g/L NaCl, plants grew more at high BZT, but only in the uninoculated treatment. Equivalently, at 0 g/L NaCl, uninoculated plants were only similar in size to inoculated plants at high BZT (Figure 2, Table 1, see also Figure S3). When NaCl was 0.8 g/L or 10 g/L, BZT concentration did not affect plant growth (Table 1). Across treatments, quadratic rates fit better and are reported (Table 1), likely because duckweed grew nearly exponentially in good conditions and died quickly at high salt.

NaCl strongly decreased plant survival, especially at higher levels of BZT. Survival was approximately 90% in treatments receiving less than 10 g/L of NaCl. At 10 g/L of NaCl, survival steadily decreased with increasing benzotriazole from about 70% with no BZT to nearly 40%, or less than half, at 10 mg/L BZT (Figure 3, Table 2). Nonetheless, the interaction between NaCl and BZT treatments was only marginally significant (Table 2). Microbial inoculation did not ameliorate the negative effects of high salt or BZT concentrations on survival; there were no significant main or interactive effects of microbes on plant survival (Table 2). Duckweed genotypes differed slightly, but non-significantly, in performance, with BH plants surviving and growing best across treatments.

**Figure 3:**
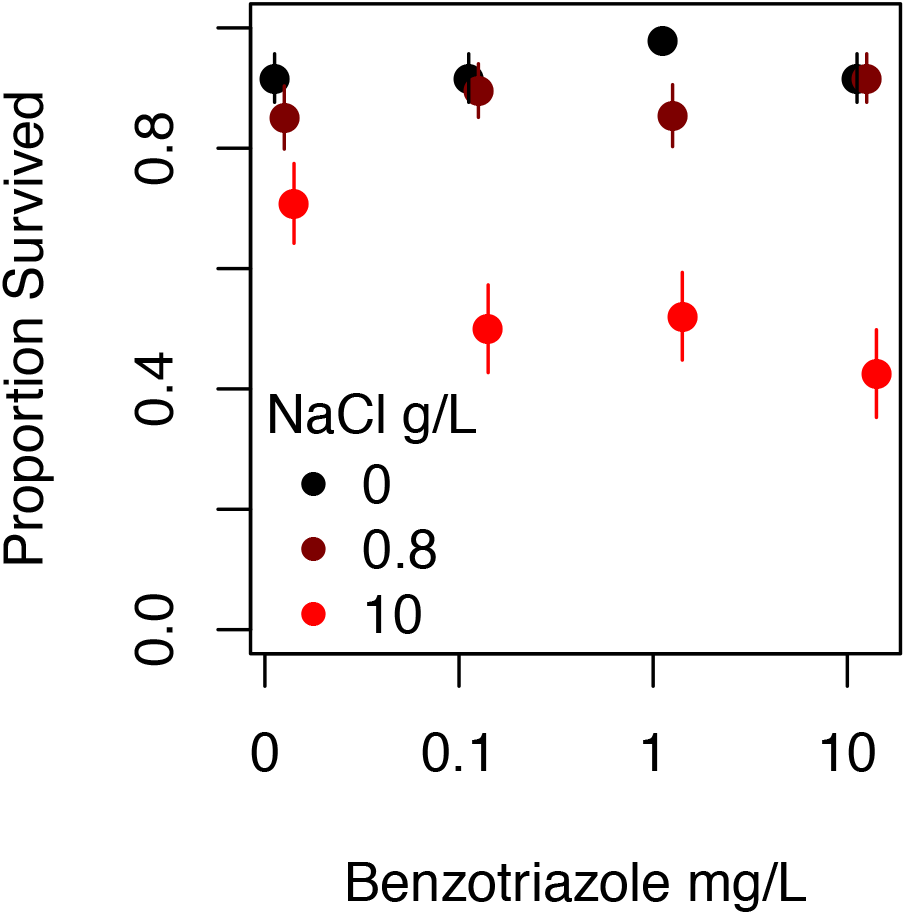
Proportion of surviving duckweed in each salt and benzotriazole treatment, with mean (points) and standard error (bars) across duckweed genotypes and inoculation treatments. Redder colors indicate higher NaCl concentrations. Points from higher salt levels are offset slightly on the x-axis to improve visualization.

**Table 2:** Model results for plant survival, plant traits, and total bacterial abundance. Significance codes:. pMCMC < 0.1, * pMCMC < 0.05, ** pMCMC < 0.01

| parameter | survival | aggregation | greenness | optical density |
| --- | --- | --- | --- | --- |
| Intercept | 103.87** | 9.61** | 0.47** | -3.90** |
| BZT | 1.18 | n.s. | n.s. | n.s. |
| NaCl | -8.79** | -0.50** | -0.0044** | -0.018** |
| Inoc | n.s. | n.s. | n.s. | NA |
| BZT:NaCl | -0.45. | n.s. | n.s. | n.s. |
| BZT:Inoc | n.s. | n.s. | n.s. | NA |
| NaCl:Inoc | n.s. | n.s. | n.s. | NA |
| BZT:Inoc:NaCl | n.s. | n.s. | n.s. | NA |

### Plant traits and microbial abundance

We found that the primary determinant of both plant traits and microbial abundance was NaCl concentration. 10 g/L of NaCl substantially reduced duckweed greenness and aggregation and total microbial abundance relative to the other treatments (Figure 4). Only minimal, non-significant differences between 0 g/L NaCl and 0.8 g/L NaCl were observed (Table 2, Figure 4). Neither benzotriazole, nor interactions between benzotriazole and other treatments, significantly affected plant greenness or aggregation or total microbial abundance (Table 2, Figure 4). Furthermore, inoculation with microbes did not affect either plant greenness or aggregation (Table 2).

**Figure 4:**
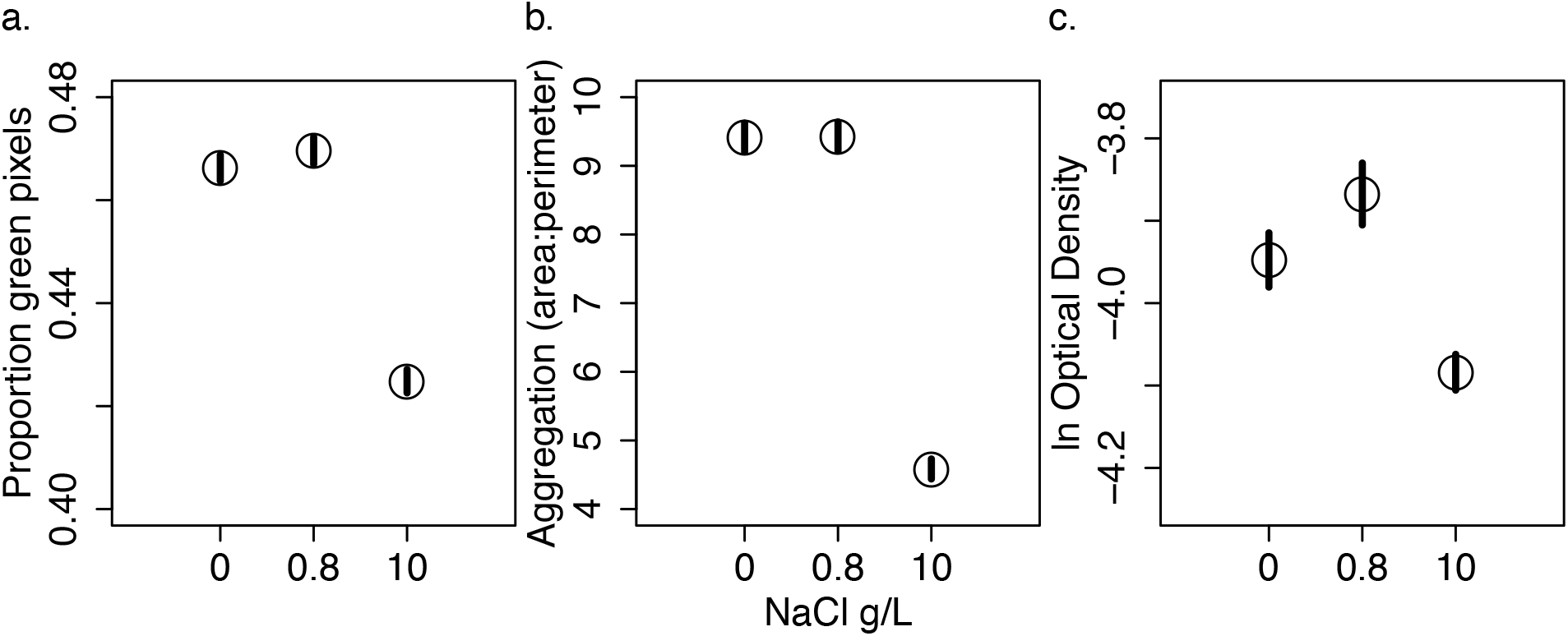
Differences among NaCl treatments in plant greenness (a), plant aggregation (b, ratio of total duckweed area and perimeter measured in pixels) and total microbial abundance (c, natural log of the optical density). Points with black bars are means and standard errors of the mean.

Nonetheless, as we expected, inoculation with microbes substantially increased optical density of well solutions at the end of the experiment, relative to uninoculated wells. The 95% HPDI for optical density in uninoculated wells was 0.014-0.015, and it was 0.018-0.020 in inoculated wells; these HDPIs were back-transformed from model predictions on a log scale and the treatment difference is statistically significant at pMCMC < 0.001.

### Benzotriazole transformation

Decreases in benzotriazole concentration from the beginning to the end of the experiment did not occur equally across treatments. BZT concentrations decreased more without salt compared to the addition of 10 g/L NaCl (pMCMC < 0.01), for lower initial benzotriazole concentrations compared to starting concentrations of 10 mg/L (pMCMC < 0.01), and in inoculated compared to uninoculated treatments (pMCMC < 0.1, see Figure 5). The highest benzotriazole concentration decreases were thus in 0 g/L NaCl, 0.1 mg/L BZT, and in the presence of microbes (64% decrease in BZT concentration for plants and microbes from BH, 58% for KSR, and 68% for MT). One exception was for BH plants and microbes at low NaCl and BZT concentrations, in which inoculated wells decreased BZT concentration less than uninoculated wells (64% and 69% respectively). While BZT generally decreased more with MT plants and microbes, and least with KSR plants and microbes, fitted random effects were not significantly different, suggesting that these *L. minor* genotypes and microbiomes have similar effects on benzotriazole transformation. Furthermore, the proportion of benzotriazole remaining was negatively correlated with duckweed final size and total microbial abundance as measured by optical density (pMCMC < 0.01 and 0.05, with each singly, respectively, < 0.05 and < 0.1 when both included in the model, see Figure 5).

**Figure 5:**
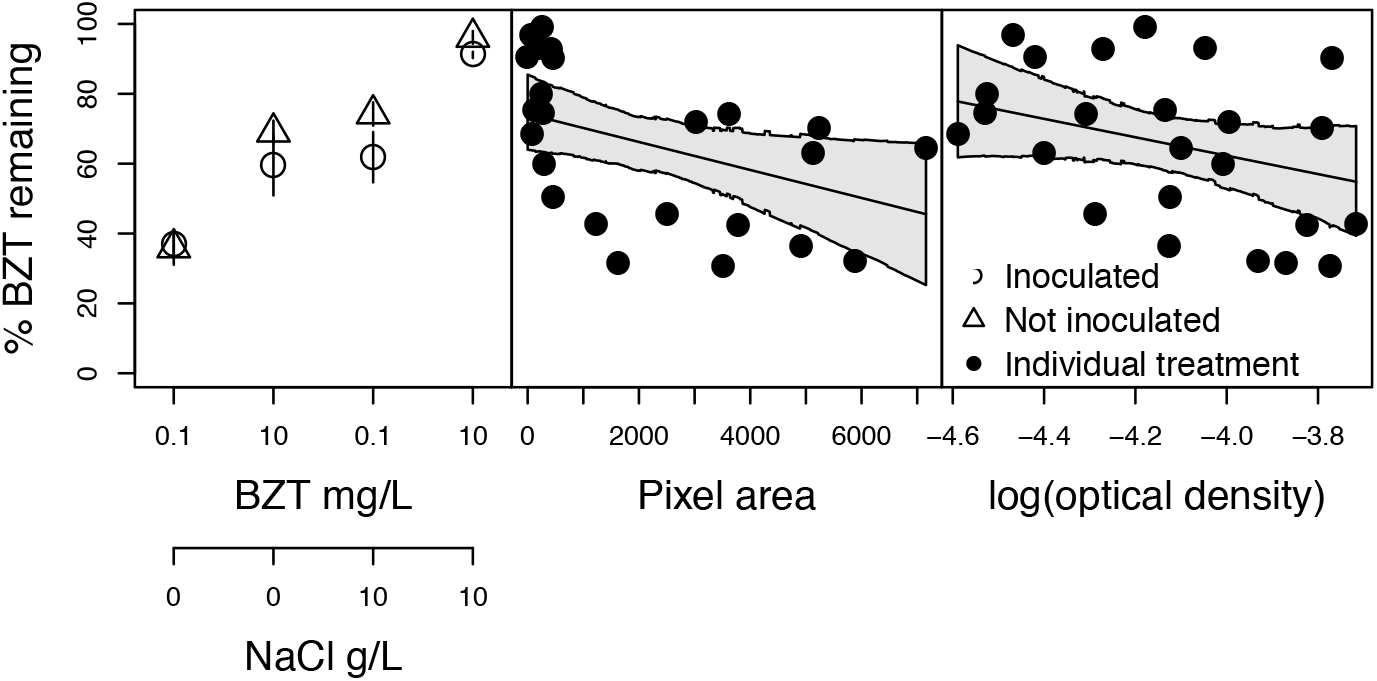
Final concentrations of BZT at the end of the experiment, as a proportion of starting concentrations. Left-most plot shows averages across BZT treatment (starting mg/L) on the x-axis, with a second x-axis below showing the corresponding NaCl treatment in g/L. Open triangles and circles indicate treatment averages for uninoculated and inoculated pooled samples, respectively. Middle and right plots show correlations with plant growth and total microbial growth, respectively, and when the other term is included in the model (solid points are raw values, not residuals).

## DISCUSSION

*Lemna minor* is an abundant and widely distributed aquatic plant, and both our results and other studies (e.g., Ishizawa et al. 2017) have found that it associates with a diverse and mostly beneficial microbial community. We experimentally manipulated these microbes as well as two common aquatic contaminants in the freshwater habitat of duckweed, sodium chloride and benzotriazole, which are both decay recalcitrant and frequently enter streams in a correlated fashion during cold season thaws (Parajulee et al., 2017). We found that the highest salt concentration, 10 g/L NaCl, was clearly stressful for duckweed and microbes; it reduced plant growth and survival, as well as made fronds less green and aggregated, while also decreasing microbial abundance (Figures 2, S1). In contrast, BZT was only stressful to plants in combination with high salt, and it never affected microbial abundance, though both plants and microbes may have contributed to the biotransformation of BZT that we observed at all but the highest levels of salt and BZT (Figures 3 5, Tables 1, 2). As in other mutualisms (e.g. Hoeksema et al., 2010), microbes had context-dependent effects on plants, with microbial inoculation increasing plant growth in most, but not all, of the environmental conditions we tested. At intermediate salt concentrations (0.8 g/L), microbes were always beneficial, but at zero salt, microbes were most beneficial at low BZT, and neutral at higher BZT. At high salt (10 g/L), microbes were mildly parasitic, although this affected only the rate of plant tissue death (Table 1) but not final plant size (Figure 2). Taken together, our results show that the effects of both microbes and contaminants on duckweed performance vary across environments, and thus that the interactive effects of microbial partners and multiple stressors may be difficult to predict in a changing world.

The three duckweed populations that we sampled associated with similar bacterial families (Figure 1), but the 16S sequencing recovered many unique bacteria at lower taxonomic levels, suggesting high turnover of bacteria across sites. We recovered taxonomic groups known to occur in abundance in the field at both coarse and fine taxonomic levels (i.e. Gammaproteobacteria, *Aeromonas*, etc.; see Figure 1 and Kivanc et al., 2011; Saxton et al., 2016). We expected that the bacteria we cultured in the lab would be a non-random subset of those that associate with wild duckweeds, and this is indeed what we found. It is well known that many microbes do not grow well under standard lab culture conditions (Rappé and Giovannoni, 2003), including specific classes of bacteria that we see in our field samples (e.g. Verrucomicrobia, Sangwan et al., 2005), limiting our ability to fully recreate natural microbiomes in experiments. Furthermore, we probably did not sequence all the microbial taxa that occurred in our field samples and cultures, as other microbes such as fungi and diatoms associate with duckweed in the field (Rejmankova et al., 1986; Goldsborough, 1993) and may also proliferate in lab culture, but are not captured by our 16S rDNA amplicon sequencing using primers optimized for bacteria. Nonetheless, our design is an improvement over many plant-microbe studies that pair taxa that may not actually associate in nature, which can drastically alter conclusions (Petipas et al., 2017). For example, meta-analyses of legume-rhizobium and plant-mycorrhizae experiments (e.g., Friesen 2012; Rúa et al. 2016) often report a preponderance of studies involving partners not known to interact in the wild. Here, we paired plants and microbes that are at least potentially locally co-adapted, having been cultured from the same field samples, even if the cultured inocula were somewhat taxonomically depauperate relative to field microbiomes.

In the plant ecology literature, the Stress Gradient Hypothesis (Callaway et al., 2002) suggests that species interactions become more beneficial under stress, but we found the opposite: microbes were more beneficial at low stress, but were neutral or slightly parasitic at the worst environmental conditions. Our results may instead be consistent with greater benefits of interactions at intermediately stressful conditions (Michalet et al., 2006; Holmgren and Scheffer, 2010); at some point a stress may become so extreme that interactions cannot ameliorate it, perhaps explaining why microbes were not beneficial at the highest salt stress. Indeed, we found slightly greater benefits of microbes at low salt compared to no stress (Table 1), and it is possible that had we explored more intermediate salt concentrations, we would have seen stronger benefits. Our study system shows promise for measuring the outcome of plant-microbe interactions across full environmental gradients, and not just at the extremes, as is often done with larger plants. Nonetheless, in the current study, there was little indication that microbes buffer plants against the negative effects of aquatic contaminants, even though we found some evidence that microbes may help degrade BZT (see below).

Our experiment found some evidence that multiple stressors have non-independent effects. If we had tested the effects of BZT concentration alone on duckweed growth, we would have come to substantially different conclusions as to whether or not it is a stressor. Here, we find that without NaCl, BZT is not stressful to plants or microbes at the concentrations we tested; in the absence of salt, there were no BZT effects on microbial growth or plant survival, and there were conditional benefits to plant growth when duckweeds were not inoculated with microbes. However, when NaCl concentrations were higher, increasing BZT concentration reduced duckweed survival, albeit with only marginal statistical significance. Thus, BZT may sometimes have negative effects on duckweed in the field, as BZT inputs to watersheds in the GTA track salt inputs nearly perfectly (Parajulee et al., 2017). Key periods of susceptibility to inputs could be during fall and spring thaw events when both concentrations are high. Ponds near snow storage sites (Alvey et al., 2016), or sites of high direct inputs, such as airfields (Breedveld et al., 2003) or dense road areas (Kiss and Fries, 2012; Parajulee et al., 2017) may be likely to reach high concentrations, especially if they have little flow through, since benzotriazole may remain in the environment for long periods of time (Breedveld et al., 2003).

Other studies have also found that high salt is detrimental to duckweed (Sree et al., 2015; Liu et al., 2017), but did not report any recovery, which we saw between days 7 and 10 (see Figure S3). This may be because we ran our experiment for longer than these previous studies. Microbial communities can shift ecololgically (Lau and Lennon, 2012; Panke-Buisse et al., 2015) or evolutionarily (Polz et al., 2013) in response to stress, and can contribute to acquired stress tolerance in plants (Lau and Lennon, 2012). However, we saw no differences in stress tolerance between uninoculated and inoculated plants, suggesting that changes occur within the duckweed themselves. Plastic developmental changes to tolerate salt may be possible for emerging daughter fronds but not adult fronds, and thus may take a while to observe under the very slow growth rates of duckweed at high salt. Indeed, transgenerational plasticity has previously been implicated in pollution tolerance (Marshall, 2008). Rapid responses are potentially critical when organisms are faced with high stress environments, and plastic development may help organisms acclimate to global change (Charmantier et al., 2008; Gimeno et al., 2009), survive in the face of pollutants (Marshall, 2008; Whitehead et al., 2010), or allow time for adaptation to stress (Whitehead et al., 2010; Ashander et al., 2016).

We expected to see toxicity of BZT for both plants and microbes, as effects have been previously detected on different microbes (Jia et al., 2006) and other *L.minor* genotypes at BZT concentrations of 10 mg/L (Seeland et al., 2012), although Gatidou et al. (2017) also found minimal impacts of 0.2 to 2 mg/L BZT on duckweed. The lack of benzotria-zole effects at lower salt and lower initial BZT concentrations may be explained by its loss during the experiment, likely through three main pathways: phototransformation, microbial transformation, or plant transformation. Even though adsorption of benzotriazole can occur, in particular at high salt concentration, it remains typically small (Rhodes-Dicker and Passeport, 2019). Benzotriazole light absorption maximum is typically around 190 to 302 nm (Borowska et al., 2016). The plastic material of the wells (polystyrene) and plastic cover sheet (polyurethane) used in the experiments filtered light wavelengths below 300 nm, thus significantly limiting the potential for phototransformation. Indeed, polyurethane’s transmittance at wavelengths lower than 300 nm is below 15%, corresponding to very large light absorption, > 0.82 absorption units (Meera et al., 2014). In addition, polystyrene’s maximum light absorption is at approximately 262 nm and it does not absorb light above 300 nm (Li et al., 1991). The absence of significant benzotriazole phototransformation was confirmed by the relatively constant benzotriazole concentrations observed in the microbe-free and high salt samples, regardless of genotype. This experimental setup therefore allowed us to specifically evaluate the role of plants and microbes for benzotriazole transformation.

The decreasing benzotriazole concentrations observed in the salt-free samples, in which duckweed growth was significantly higher than in the corresponding samples with 10 g/L NaCl, suggest a biological process. Since this occurred even in the uninoculated treatments, it suggests that duckweed plants played a role in benzotriazole removal, as has been suggested previously (Gatidou et al., 2017). In other systems, microbes can increase phytodegradation simply by positively impacting plant growth rate (Glick, 2003; Sobariu et al., 2017). Microbes can also alter the rates at which plants bioaccumulate contaminants (Burd et al., 2000), or they can directly metabolize compounds themselves (McGuinness and Dowling, 2009). In most cases, inoculation with microbes resulted in even lower final concentrations, even after accounting for duckweed growth, final microbial abundance was negatively correlated with the proportion of remaining benzotriazole (Figure 5), suggesting a role for microbes beyond impacts on plant growth. In fact, similar microbes inhabiting the roots of other duckweed species have been shown to directly impact degradation of other recalcitrant environmental contaminants, including phenol (Toyama et al., 2009) and 4-tert-Butylphenol (Ogata et al., 2013). Furthermore, some microbial digestion of benzotriazole has been shown to occur in different systems, primarily waste water treatment plant sludge (Gruden et al., 2001; Weiss et al., 2006), which can include microbes from the same taxa we observe here. Of note, ASVs identified as *Pseudomonas veronii*, a taxon implicated in biotransformation of other aromatic compounds (e.g., Nam et al., 2003) comprised 18% of sequences from MT, the source that resulted in the most BZT biotransformation here.

We note that the BZT loss rates we observe here are slightly lower than those observed previously for *L. minor* (Gatidou et al., 2017). Lower light conditions in our experiment may have reduced plant growth and therefore reduced transformation rates, or variation may exist across duckweed genotypes in transformation capacity, as it does differ across species (LeFevre et al., 2017). However, the largest differences may come from our inclusion of NaCl treatments and higher starting levels of benzotriazole, which both increased the final concentrations of benzotriazole (Figure 5). Environments highly co-contaminated with benzotriazole and salt may reduce the transformation of benzotriazole by duckweed-microbe systems, but benzotriazole concentrations in the environment are typically closer to 0.1 than 10 mg/L (Loos et al., 2010), and even though benzotriazole and salt are both detected at higher concentrations in winter (Parajulee et al., 2017), NaCl levels of 10 g/L are less frequent than 0.8 g/L (Rhodes-Dicker and Passeport, 2019). Thus, these results suggest that high benzotriazole removal can still be expected in most shallow benzotriazole-contaminated and duckweed-populated waters.

## CONCLUSIONS

Stressors can shift outcomes of species interactions, sometimes strengthening positive outcomes (He et al., 2013), and sometimes disrupting them (Shantz et al., 2016), yet we have relatively few tests of the effects of multiple novel stressors on species interactions. We find here that under simultaneous stress from two common aquatic contaminants, a widespread plant-microbe mutualism neither exacerbated nor mitigated the negative effects of stress, counter to results for nutrient pollution (Shantz et al., 2016). We still know very little about the non-additive effects of the many concurrent global changes presently impacting species interactions, but duckweeds and their microbiota offer a tractable experimental system for rapid and highly multifactorial manipulations of abiotic and biotic environments.

## Acknowledgements

The authors would like to thank E. Lash for help collecting duckweeds, and students and volunteers who have contributed to maintaining duckweed and microbe cultures in the lab.

## Author contributions

All authors contributed substantially to the design of the study, provisioning of materials, or revising of the manuscript. AO, MEF, EP proposed the study; EP, MEF, and JL provisioned materials; AO, ZY, and DL collected the data; AO and ZY performed analyses; AO and MEF provided the first draft of the manuscript.

The project was funded by NSERC Discovery Grants and a University of Toronto XSeed Grant to MEF and EP.

## Data Accessibility Statement

Upon publication data will be accessible via figshare. Scripts will be accessible on GitHub.

## Appendix S1

**Table S1:** Locations of source sites for duckweed and associated microbes.

| Site | Latitude | Longitude | Type of site |
| --- | --- | --- | --- |
| Koffler Science Reserve | 44.0350 | -79.5178 | Rural reserve |
| Bruce Handscombe Memorial Park | 43.8172 | -79.0975 | Suburban park |
| Moccasin Trail Park | 43.7333 | -79.3319 | Urban park |

**Table S2:**
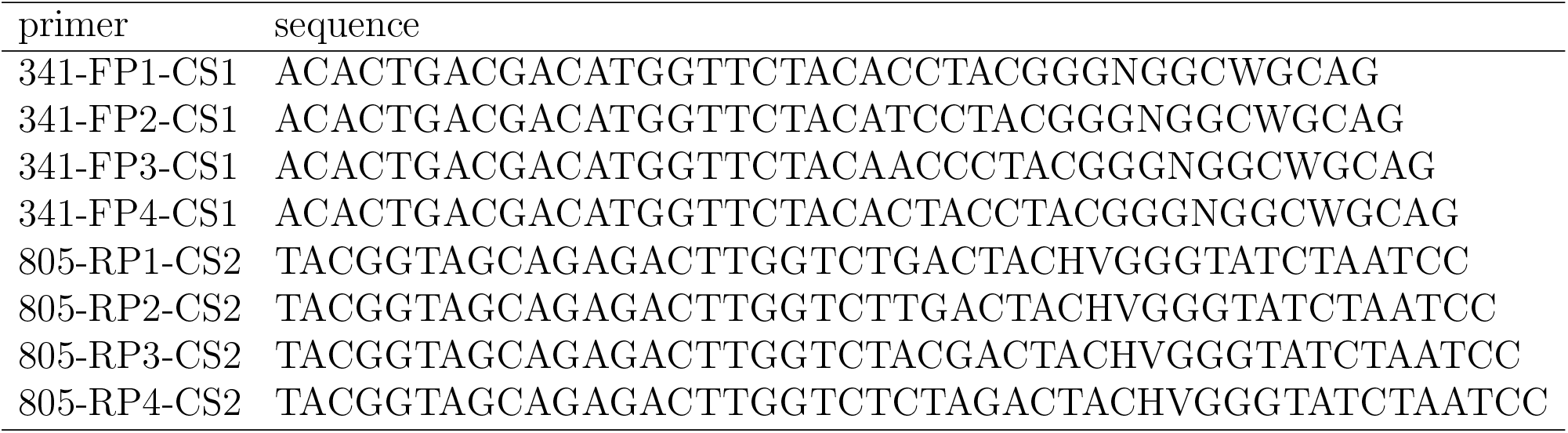
Primer sequences for 16s rDNA V3-V4 region used.

**Figure S1:**
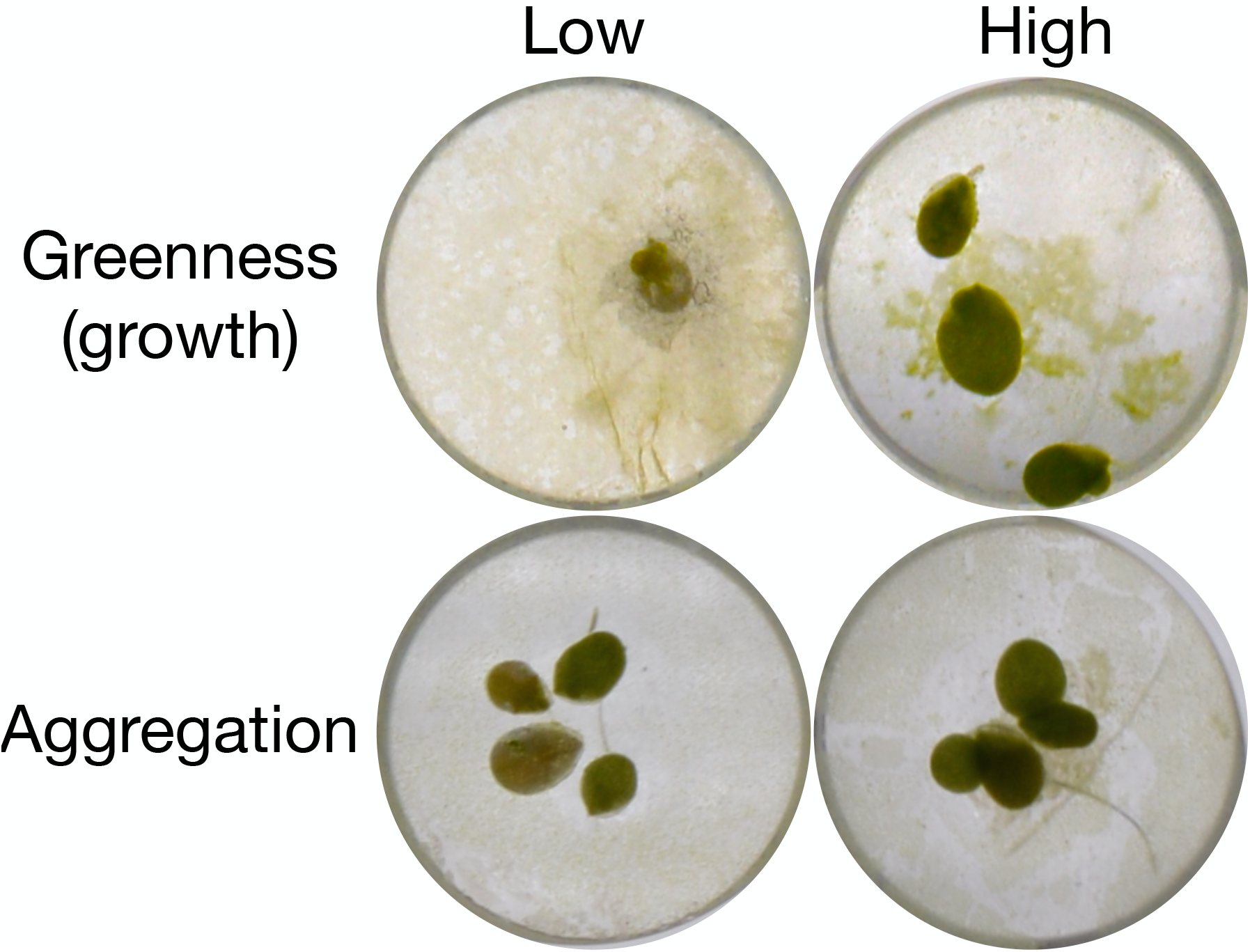
Example duckweed in wells of plates showing lower (left images) and higher (right images) greenness (top row) and aggregation (bottom row), as photographed at the end of the experiment. The top row images additionally show smaller (left) and larger (right) pixel area.

**Figure S2:**
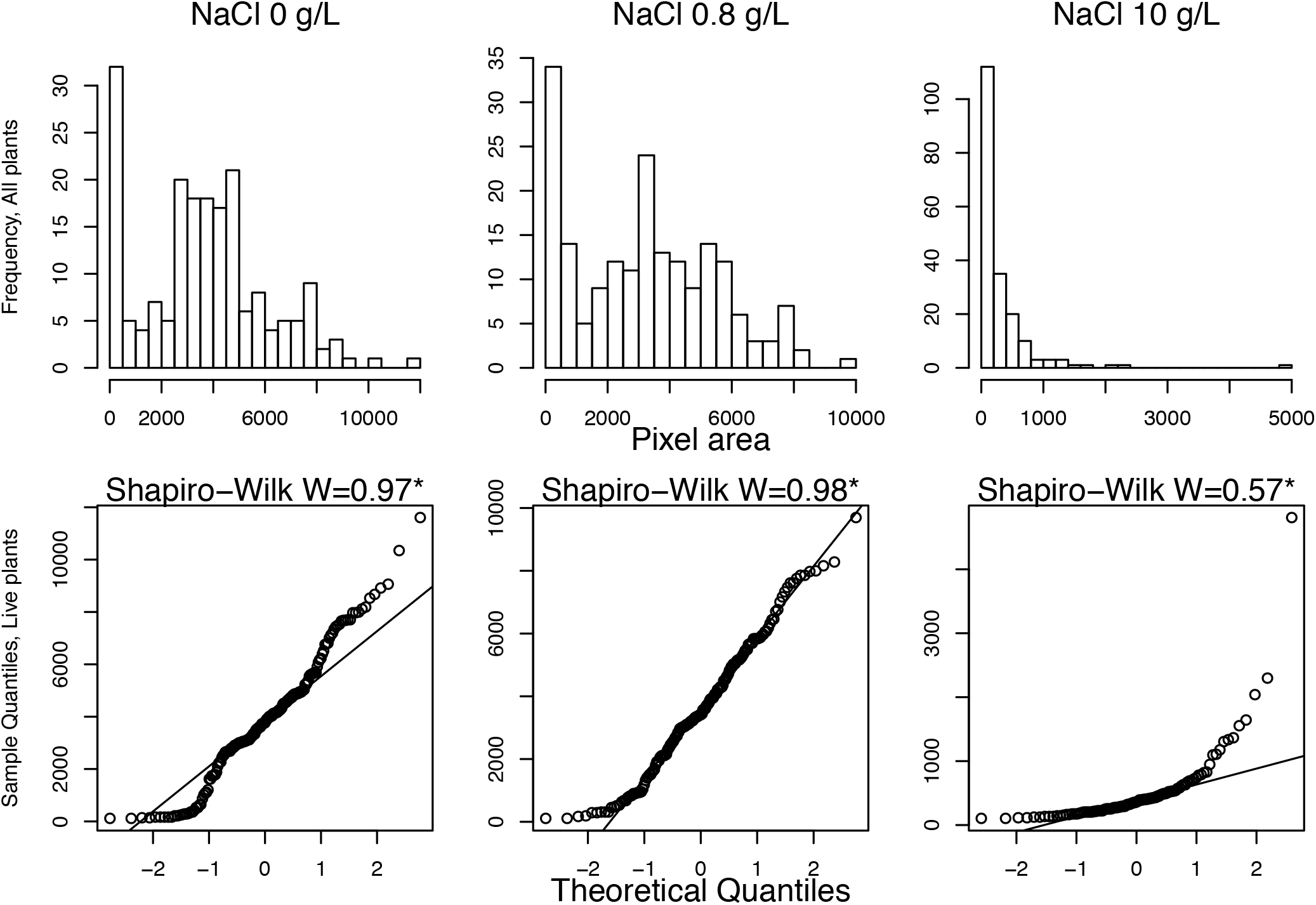
Normality tests for pixel area data. Only data from the final 10-day time point is shown here. Day 10 is the least normally distributed due to the most accumulated dead plants. Histograms of data from all levels of salt clearly show an excess of zeros. When excess zeros are removed (i.e., when dead plants are excluded), only high salt treatment data deviate strongly from normality expectations in the quantile-quantile plots. *Shapiro-Wilk test significant for all salt treatments, but trivial deviation from W=1 for lower salt treatments.

**Figure S3:**
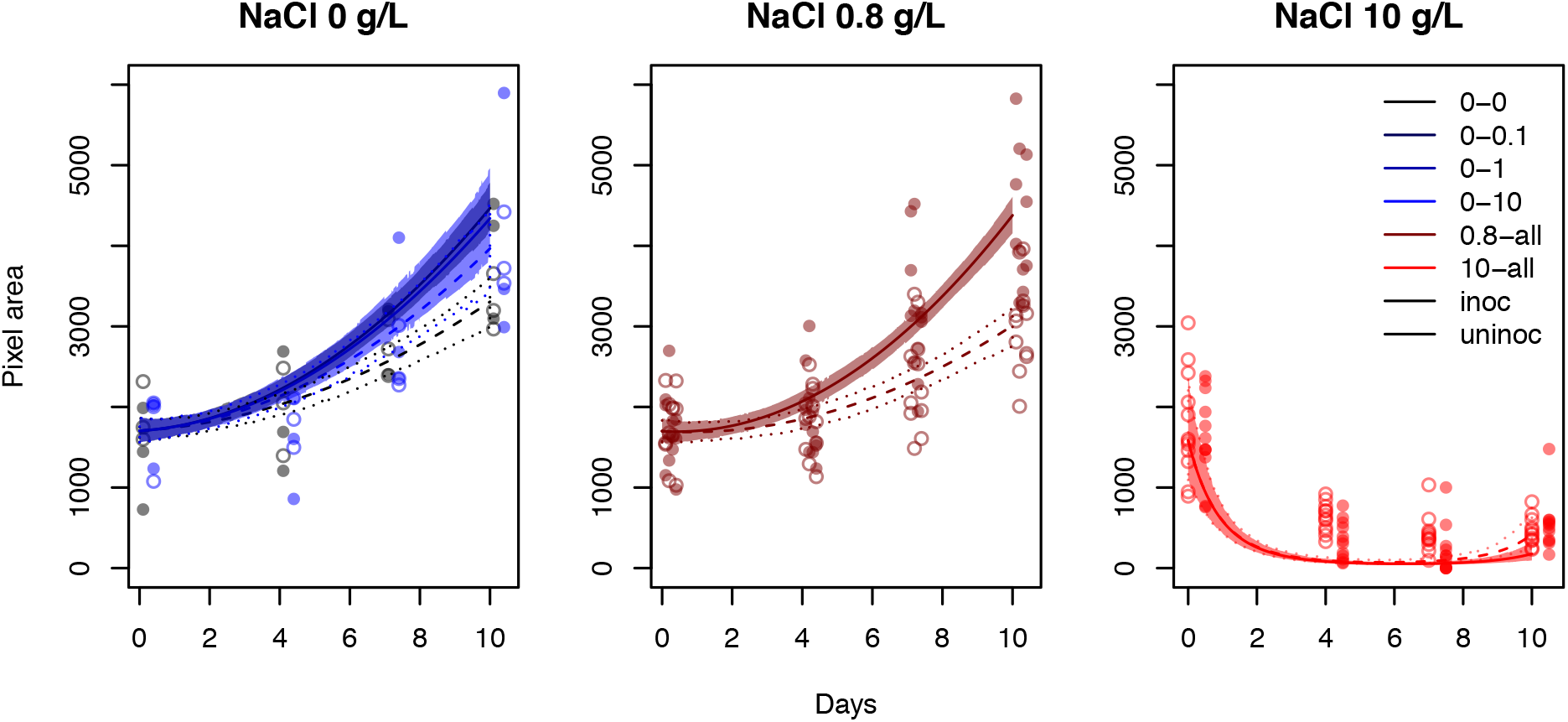
Growth in pixel area of surviving duckweeds over time. Each NaCl treatment was analyzed separately, using Gaussian (0 and 0.8 g/L NaCl) or Poisson (10 g/L NaCL) distributions. Experimental treatment means (points with standard error bars; offset on the x-axis for inoculated plants) and model predictions (predicted mean lines and 95% HPDIs) shown separately for all fixed effects significant in models (at 10 and 0.8 g/L salt only inoculated and uninoculated treatments are shown separately). Filled circles, and filled intervals with solid lines indicate inoculated plant data and predictions, respectively. Open circles, and open intervals with dashed lines indicate uninoculated plant data and predictions, respectively. Few differences exist across levels of BZT, even at 0 g/L salt where they are significant, so we show only intervals for the highest and lowest BZT level, and only for 0 g/L salt.

